# Strong SARS-CoV-2 N-specific CD8^+^ T immunity induced by engineered extracellular vesicles associates with protection from lethal infection in mice

**DOI:** 10.1101/2022.01.10.475620

**Authors:** Flavia Ferrantelli, Chiara Chiozzini, Francesco Manfredi, Patrizia Leone, Massimo Spada, Antonio Di Virgilio, Andrea Giovannelli, Massimo Sanchez, Andrea Cara, Zuleika Michelini, Maurizio Federico

## Abstract

SARS-CoV-2-specific CD8^+^ T cell immunity is expected to counteract viral variants in both efficient and durable ways. We recently described a way to induce a potent SARS-CoV-2 CD8^+^ T immune response through the generation of engineered extracellular vesicles (EVs) emerging from muscle cells. This method relies on intramuscular injection of DNA vectors expressing different SARS-CoV-2 antigens fused at their N-terminus with Nef^mut^ protein, i.e., a very efficient EV-anchoring protein. However, quality, tissue distribution, and efficacy of these SARS-CoV-2-specific CD8^+^ T cells remained uninvestigated. To fill the gaps, antigen-specific CD8^+^ T lymphocytes induced by the immunization through the Nef^mut^-based method were characterized in terms of their polyfunctionality and localization at lung airways, i.e., the primary targets of SARS-CoV-2 infection. We found that injection of vectors expressing Nef^mut^/S1 and Nef^mut^/N generated polyfunctional CD8^+^ T lymphocytes in both spleens and bronchoalveolar lavage fluids (BALFs). When immunized mice were infected with 4.4 lethal doses 50% of SARS-CoV-2, all S1-immunized mice succumbed, whereas those developing the highest percentages of N-specific CD8^+^ T lymphocytes resisted the lethal challenge. We also provide evidence that the N-specific immunization coupled with the development of antigen-specific CD8^+^ T-resident memory cells in lungs, supporting the idea that the Nef^mut^- based immunization can confer a long-lasting, lung-specific immune memory. In view of the limitations of current anti-SARS-CoV-2 vaccines in terms of antibody waning and efficiency against variants, our CD8^+^ T cell-based platform could be considered for a new combination prophylactic strategy.

## INTRODUCTION

The term “correlate of protection” refers to a laboratory parameter associated with protection from a clinical disease [1].The immunological correlates of protection against SARS-CoV-2 infection have not been identified yet. A coordinated action of innate immunity, CD4^+^ T cells, CD8^+^ T cells, and neutralizing antibodies is most likely necessary for natural control SARS-CoV-2 infection. In this scenario, neutralizing antibodies certainly play a key role in protecting from infection, and, accordingly, vaccine strategies aimed at producing anti-SARS-CoV-2 neutralizing antibodies are now available. Those based on the administration of lipidic nanovesicle-complexed messenger (m)RNA molecules appear to be the most effective. However, their overall efficiency is challenged by emergence of variants and antibody waning.

The role of CD8^+^ T cell immunity in the recovery from SARS-CoV-2 infection has been widely demonstrated. For instance, a seminal study on rhesus macaques demonstrated that the depletion of CD8^+^ T cells after a first virus challenge abolished the protective effect of natural immunity against a virus re-challenge applied after waning of neutralizing antibodies [2]. The authors concluded that the antiviral CD8^+^ T cell immunity can control viral spread in the context of suboptimal levels of neutralizing antibodies. In humans, the presence of virus-specific CD8^+^ T lymphocytes associates with a better recovery from the disease [3-6]. SARS-CoV-2-specific CD8^+^ T cells develop also in recovered individuals who did not produce anti-SARS-CoV-2 antibodies [7]. Of major relevance, SARS-CoV-2 specific CD8^+^ T cell immunity maintains intact its efficacy in the presence of the amino acid substitutions occurring in the Spike (S) protein of emerging viral variants [8-10]. In addition, SARS-CoV survivors preserved N-specific CD8^+^ T lymphocytes for as many as 16 years after recovery [11].

Up until now, no vaccine strategy specifically devoted to the induction of CD8^+^ T immunity has been proposed for humans. To counteract SARS-CoV-2 infection, we applied an original CD8^+^ T cell-based vaccine platform previously proven to be effective against both HPV16-and HER2-induced cancers [12, 13]. This method was conceived to induce antigen-specific cytotoxic CD8^+^ T lymphocyte (CTL) immunity, and is based on *in vivo* engineering of extracellular vesicles (EVs).

All cell types constitutively release different types of nanovesicles which are collectively referred to as EVs [14]. Our vaccine platform is based on intramuscular (i.m.) injection of DNA vectors coding for Nef^mut^, i.e., a biologically inactive Human Immunodeficiency Virus (HIV)-Type 1 Nef protein. This protein mutant shows an extraordinarily high efficiency of incorporation into EVs even when foreign polypeptides are fused to its C-terminus [15, 16]. Intracellular expression of Nef^mut^-derivatives leads to their incorporation into EVs physiologically released by host cells. After i.m. injection of Nef^mut^-based vectors, these engineered nanovesicles are released by muscle cells, can freely circulate into the body and be internalized by antigen-presenting cells (APCs). EV-associated antigens are then cross-presented to prime antigen-specific CD8^+^ T-lymphocytes [17].

Injection of DNA vectors expressing the products of fusion between Nef^mut^ and either SARS-CoV-2 S1 or N was shown to generate strong antigen-specific CD8^+^ T cell immunity in spleen as detected by IFN-γ EliSpot analysis [18]. Here, we expanded the investigations towards the functional characterization of virus-specific CD8^+^ T cells in both spleen and lungs. In addition, their antiviral efficacy was tested by virus challenge experiments infecting immunized transgenic mice with a lethal dose of SARS-CoV-2. We found that the generation of high levels of nucleocapsid (N)-specific polyfunctional CD8^+^ T lymphocytes in both spleen and lungs associated with resistance to the lethal effect of challenging virus.

## MATERIALS AND METHODS

### DNA constructs

Open-reading frames coding for Nef^mut^ fused with either S1, S2, or N SARS-CoV-2 proteins were cloned into pVAX1 plasmid (Thermo Fisher) as previously described [18]. In these constructs, S1 spans from aa 19, i.e., downstream of the signal peptide, to aa 680, just upstream of the furin-like cleavage site; S2 included the extracellular portion of the subunit with the exclusion of the two fusion domains; finally, the entire N protein (422 aa), except M1 amino acid, was fused to Nef^mut^. A GPGP linker was inserted between Nef^mut^ and downstream sequences. Stop codons of SARS-CoV-2-related sequences were preceded by sequences coding for a DYKDDDK epitope tag (flag-tag). SARS-CoV-2 sequences were optimized for expression in human cells through GeneSmart software from Genescript. All vectors were synthesized by Explora Biotech.

### Animals and authorizations

Six-weeks old C57 Bl/6 and C57 Bl/6 K18**-**hACE-2 transgenic [19] female mice were purchased from Charles River and hosted at the Central and BSL3 Animal Facilities of the Istituto Superiore di Sanità, as approved by the Italian Ministry of Health, authorizations 565/2020 and 591/2021 released on June 3^rd^ 2020 and July 30^th^ 2021, respectively. Before the first procedure, DATAMARS microchips were inserted sub-cute on the dorsal midline between the shoulder blades.

### Mouse immunization

Isoflurane-anesthetized mice were inoculated i.m. with 10 μg of DNA in 30 μL of sterile, 0.9% saline solution. DNA injection was immediately followed by electroporation at the site of inoculation, with an Agilpulse BTX device, using a 4-needle electrode array (4 mm gap, 5 mm needle length) and applying the following parameters: 1 pulse of 450 V for 50 μs; 0.2 ms interval; 1 pulse of 450 V for 50 μs; 50 ms interval; 8 pulses of 110 V for 10 ms with 20 ms intervals. Mice were immunized into both quadriceps, twice, 2 weeks apart. Fourteen days after the second immunization, mice were sacrificed by either cervical dislocation or CO_2_ inhalation.

### Isolation of cells from blood, bronchoalveolar lavage fluids (BALFs), lungs, and spleen

For pre-infection immunity assessment, mice were bled by retro orbital puncture under topical anesthesia. Peripheral blood mononuclear cells (PBMCs) were recovered from EDTA-blood samples after erythrocyte removal by treatment with ACK lysing buffer (Gibco) according to the manufacturer’s instructions.

To conduct bronchoalveolar lavages, CO_2_-sacrificed mice were laid on their back, neck skin was cut open along the median line, and muscles were open apart to expose the trachea. A piece of surgical thread was tied around the trachea and a 1-millimeter cut was performed between two cartilage rings, to insert a 22 G Exel Safelet catheter 0.5 cm down into the trachea. Before proceeding with lavage, the catheter was firmly tied into the trachea by the surgical thread. An insulin syringe was loaded with 1 mL of cold 1× PBS, attached to the catheter and used to gently inject the buffer into the lungs and back, twice, while massaging the mouse thorax. Bronchoalveolar lavage fluid (BALF) was placed on ice into a 15 mL conical tube, and lavage repeated two more times to recover a total lavage volume averaging 2.5 mL/mouse.

Mouse lungs were excised, extensively washed with 1× PBS, cut into small pieces and then digested for 30 minutes under gentle agitation at 37 °C with 7 mL of the following solution: type III collagenase (Worthington Biochemical), 4 mg/mL, DNase I (Sigma), 0.05 mg/mL, resuspendend in 1×PBS. After digestion, an equal volume of medium was added, the extract was passed through a 70 μm cell strainer, washed, and recovered cells resuspended in 1×PBS/ACK (1:1 v/v) for red blood cells lysis.

Spleens were explanted, placed into tubes containing 1 mL of RPMI 1640 and 50 μM 2-mercaptoethanol, then transferred into 60 mm Petri dishes with 2 mL of the same medium. Splenocytes were obtained by notching the spleen sac and pushing the cells out with the plunger seal of a 1 mL syringe. After addition of 2 mL of medium, cells were transferred into a 15 mL conical tube, and the Petri plate washed with 4 mL of medium to maximize cell recovery. Afterwards, cells were collected by centrifugation, resuspended in RPMI complete medium containing 50 μM 2-mercaptoethanol and 10% heat-inactivated fetal calf serum (FCS), and counted before being used in IFN-γ EliSpot and/or intracellular cytokine staining (ICS) assays.

### IFN-γ EliSpot assay

A total of 2.5×10^5^ live cells were seeded in triplicate in microwells of 96-multiwell plates (Millipore) previously coated with the anti-mouse IFN-γ AN18 mAb (Mabtech) in RPMI 1640, 10% FCS, 50 μM 2-mercaptoethanol. Cell cultures were carried out for 16 h in the presence of 5 μg/mL of CD8-specific SARS-CoV-2 H2-b peptides S1: 539–546 VNFNFNGL [20], and N: 219–228 ALALLLLDRL [21]. As negative controls, 5 μg/mL of H2-b binding peptides were used. Peptide preparations were obtained from BEI resources. To check for cell responsiveness, 10 ng/mL phorbol 12-myristate 13-acetate (PMA, Sigma) plus 500 ng/mL of ionomycin (Sigma) were added to the cultures. After 16 h, cells were discarded, and the plate was incubated for 2 h at room temperature with R4-6A2 biotinylated anti-IFN-γ antibody (Mabtech) at the concentration of 100 μg/mL. Wells were then washed and treated for 1 h at room temperature with 1:1,000 diluted streptavidin-ALP from Mabtech. Afterwards, 100 μL/well of SigmaFast BCIP/NBT were added to the wells to develop spots. Spot-forming cells were finally analyzed and counted using an AELVIS EliSpot reader.

### Intracellular cytokine staining (ICS) and flow cytometry analysis

Cells collected from BALFs, spleens and lungs were cultured at 1 × 10^7^/mL in RPMI medium, 10% FCS, 50 μM 2-mercaptoethanol (Sigma), 1 μg/mL brefeldin A (BD Biosciences), and in the presence of 5 μg/mL of either S1, N, or unrelated H2-b CD8-specific peptides. Positive controls were conducted by adding 10 ng/mL PMA (Sigma) plus 1 μg/mL ionomycin (Sigma). After 16 h, cells were stained with 1 μL of LIVE/DEAD Fixable FVD-eFluor506 Dead Cell reagent (Invitrogen Thermo Fisher) in 1 mL of 1×PBS for 30 min at 4 °C, and excess dye removed by 2 washes with 500 μL of 1×PBS. Non-specific staining was minimized by pre-incubating cells with 0.5 μg of Fc blocking mAbs (i.e., anti-CD16/CD32 antibodies, Invitrogen/eBioscience Thermo Fisher) in 100 μL of 1×PBS with 2% FCS for 15 min at 4 °C. Staining for cell surface markers was performed upon incubation for 1 h at 4 °C with 2 μL of the following anti-mouse Abs: FITC-conjugated anti-CD3, APC-Cy7-conjugated anti-CD8a, PerCP-conjugated anti-CD4, and BUV395-conjugated anti-CD44 (BD Biosciences). CD8^+^ T-resident memory (Trm) cells were identified by staining with BUV750-conjugated anti-CD49a, PECF594-conjugated anti-CD69, and BUV563-conjugated anti-CD103 (BD Biosciences). After washing, cells were fixed and permeabilized using the Cytofix/Cytoperm kit (BD Biosciences), according to the manufacturer’s recommendations. For intracellular cytokine staining (ICS), cells were labeled for 1 h at 4 °C with 2 μL of the following Abs: PE-Cy7-conjugated anti-IFN-γ, PE-conjugated anti-IL-2 (Invitrogen/eBioscience Thermo Fisher), and BV421 rat anti-TNF-α (BD Biosciences) in a total of 100 μL of 1× Perm/Wash Buffer (BD Biosciences). After two washes, cells were fixed in 200 μL of 1× PBS/formaldehyde (2% v/v). Samples were then analyzed by a CyotFLEX LX (Beckman Coulter, Brea, CA, USA) flow cytometer and analyzed using Kaluza software (Beckman Coulter). Gating strategy was as follows (Supplementary fig.1): live cells as assessed by LIVE/DEAD Dye vs. FSC-A, singlet cells from FSC-A vs. FSC-H (singlet 1) and SSC-A vs. SSC-W (singlet 2), CD3^+^ cells from CD3-FITC vs. SSC-A, CD8^+^, or CD4^+^ cells from CD8-APC-Cy7 vs. CD4-PerCP. The CD8^+^ T cell population was gated against CD44^+^ cells, and the population of cells positive for both CD8 and CD44 was analyzed for APC-Cy7, PE, and BV421 to detect changes in IFN-γ, IL-2, and TNF-α production, respectively. Boolean gates were created to determine cytokine co-expression patterns.

### SARS-CoV-2 preparation and in vitro titration

VERO-E6 cells were grown in DMEM (Gibco) supplemented with 2% FCS, 100 units/ml penicillin, 100 μg/ml streptomycin, 2 mM L-glutamine, 1 mM sodium pyruvate, and 1× non-essential amino acids (Gibco). The ancestral viral isolate SARS-CoV-2/Italy INMI1#52284 (SARS-Related Coronavirus 2, isolate Italy-INMI1, NR-52284, deposited by Dr. Maria R. Capobianchi for distribution through BEI Resources, NIAID, NIH) was propagated by inoculation of 70% confluent VERO-E6 cells in 175 cm^2^ cell culture flasks [22]. Infected cell culture supernatant was harvested at 72 h post infection, clarified, aliquoted, and stored at -80 °C.

To determine SARS-CoV-2 stock TCID_50_ (tissue culture infectious doses 50%), 2.2 × 10^4^ Vero E6 cells/well were added onto 96-well plates (Corning, Mediatech Inc) and, the next day, octuplicate cultures were inoculated with 10-fold serial dilutions of virus (100 μL/well). Cells were incubated for 5–6 days and then checked daily for cytopathic effect.

### Mouse infection

Before experimental infection, mice were anesthetized with a combination of ketamine (50 mg/kg of body weight) and medetomidine (1 mg/kg of body weight) administered intraperitoneally (IP). After virus challenge, intraperitoneal injection of atipamezole (1 mg/kg of body weight) was used as a reversal agent.

For *in vivo* titration of the SARS-CoV-2/Italy INMI1#52284 isolate, age-matched K18-hACE-2 mice were randomized by body weight into groups of 4, and challenged with 5-fold serial dilutions in 1×PBS of the virus preparation containing 2.2×10^5^, 4.4×10^4^, 8.8×10^3^, or 1.8×10^3^ TCID_50_. A volume of 30 μL of each dilution was administered intranasally, 15 μL per nostril by slowly pipetting at a depth of 1-2 mm. As a negative control, 4 mice were sedated and an equal volume of 1×PBS was administered. Animals were assessed daily for clinical signs of infection and weight loss over a 15-day follow-up.

Virus challenge was performed similarly in immunized mice, using a virus dose of 4.4×10^4^ TCID_50_ corresponding to 4.4 lethal doses 50% (LD_50_), and resulting in 99.99% predicted probability of mortality.

### Statistical analysis

When appropriate, data are presented as mean + standard deviation (SD). Virus titer *in vitro* was determined as TCID_50_ with the Spearman-Karber method. For the *in vivo* titration, virus dilutions and number of deaths/group after challenge were used to calculate the LD_50_ (Quest Graph™ LD_50_ Calculator, AAT Bioquest, Inc.). The Kaplan-Meier curve was used to show survival rate differences among groups of animals challenged with different doses of SARS-CoV-2 virus, or immunized and infected with 4.4 LD_50_ of the same virus stock. The log-rank test was used to compare survival in different vaccination groups. When indicated, the Mann-Whitney U test was conducted. *p* < 0.05 was considered statistically significant.

## RESULTS

### Induction of polyfunctional antigen-specific CD8^+^ T lymphocytes after i.m. injection of DNA vectors expressing either SARS-CoV-2 S1 or N fused with Nef^mut^

The i.m. injection of DNA vectors expressing Nef^mut^-based fusion products leads to their incorporation into EVs spontaneously released by muscle cells [16, 23]. We previously showed that SARS-CoV-2 S1 and N antigens can be uploaded in engineered EVs, and i.m. injection of respective DNA vectors led to the induction of antigen-specific CD8^+^ T cells as revealed by IFN-γ EliSpot analysis [18]. However, quality, biodistribution, and effectiveness of such SARS-CoV-2-specific CD8^+^ T immunity remained essentially unexplored. To fill the gaps, we first analyzed the polyfunctionality of antigen-specific CD8^+^ T cells induced through the Nef^mut^-based method. To this aim, C57 Bl/6 mice were injected with DNA vectors expressing either Nef^mut^/S1, Nef^mut^/N (Fig. 1) or, as control, Nef^mut^ alone. Fifteen days after the second inoculation, splenocytes were isolated and incubated overnight with either specific or MHC Class I-matched, unrelated peptides. Through ICS/flow cytometry analysis we found 5-15% of antigen-specific cells expressing either IFN-γ, IL-2, or TNF-α within the CD8^+^/CD44^+^ subpopulations (Fig. 2A-B, and Supplementary fig. 1). The analysis of combined cytokine expression revealed the presence of as many as 25-30% triple positive cells within the activated cell populations (Fig. 2C).

**Figure 1.**
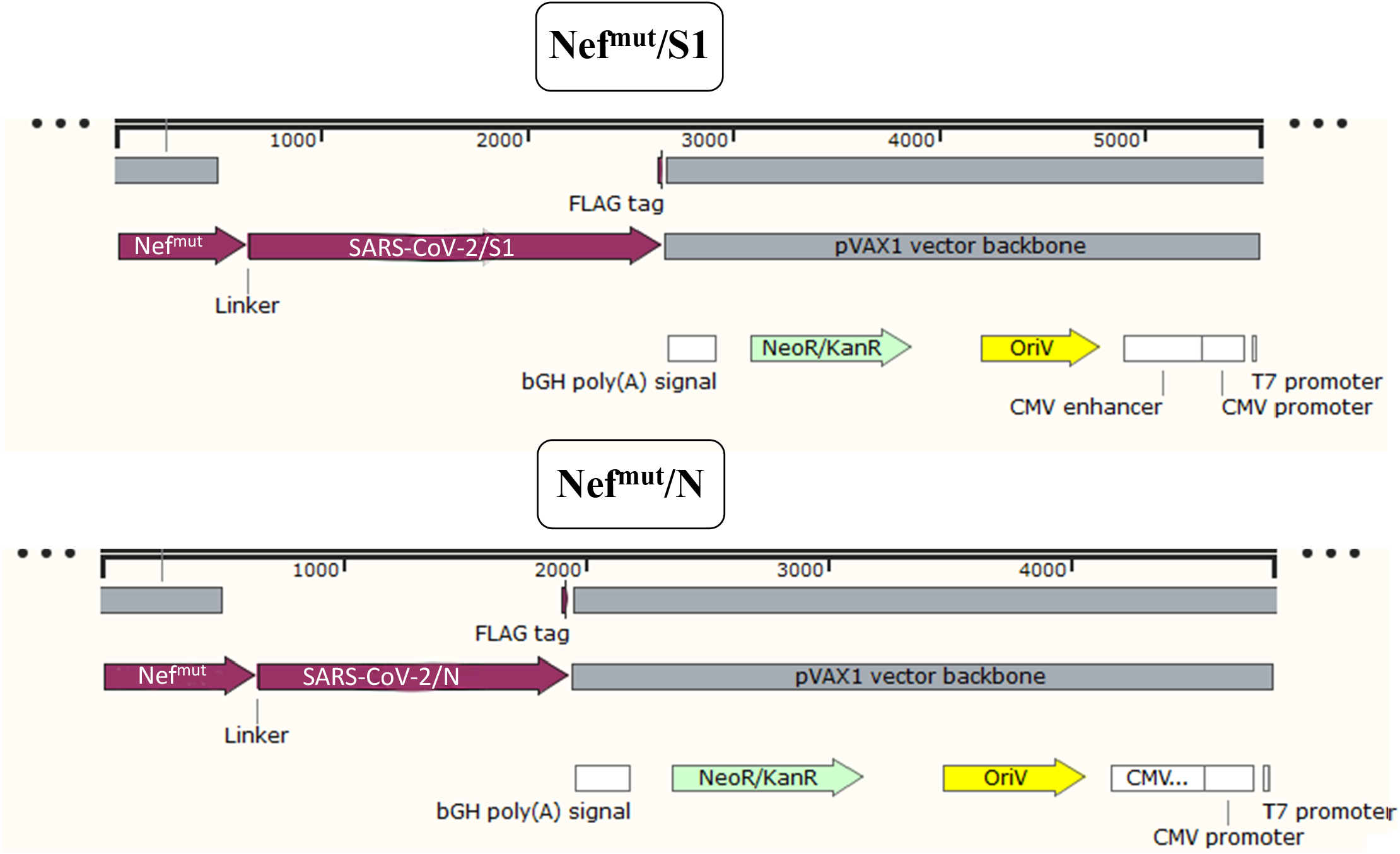
Linear maps of vectors expressing SARS-CoV-2-based fusion proteins. Shown are the structure of pVAX1 vectors expressing either S1 or N proteins fused with Nef^mut^. Positions of fusion products, functional regions of the vectors, as well as both GPGP linker and Flag-tag are indicated.

**Figure 2.**
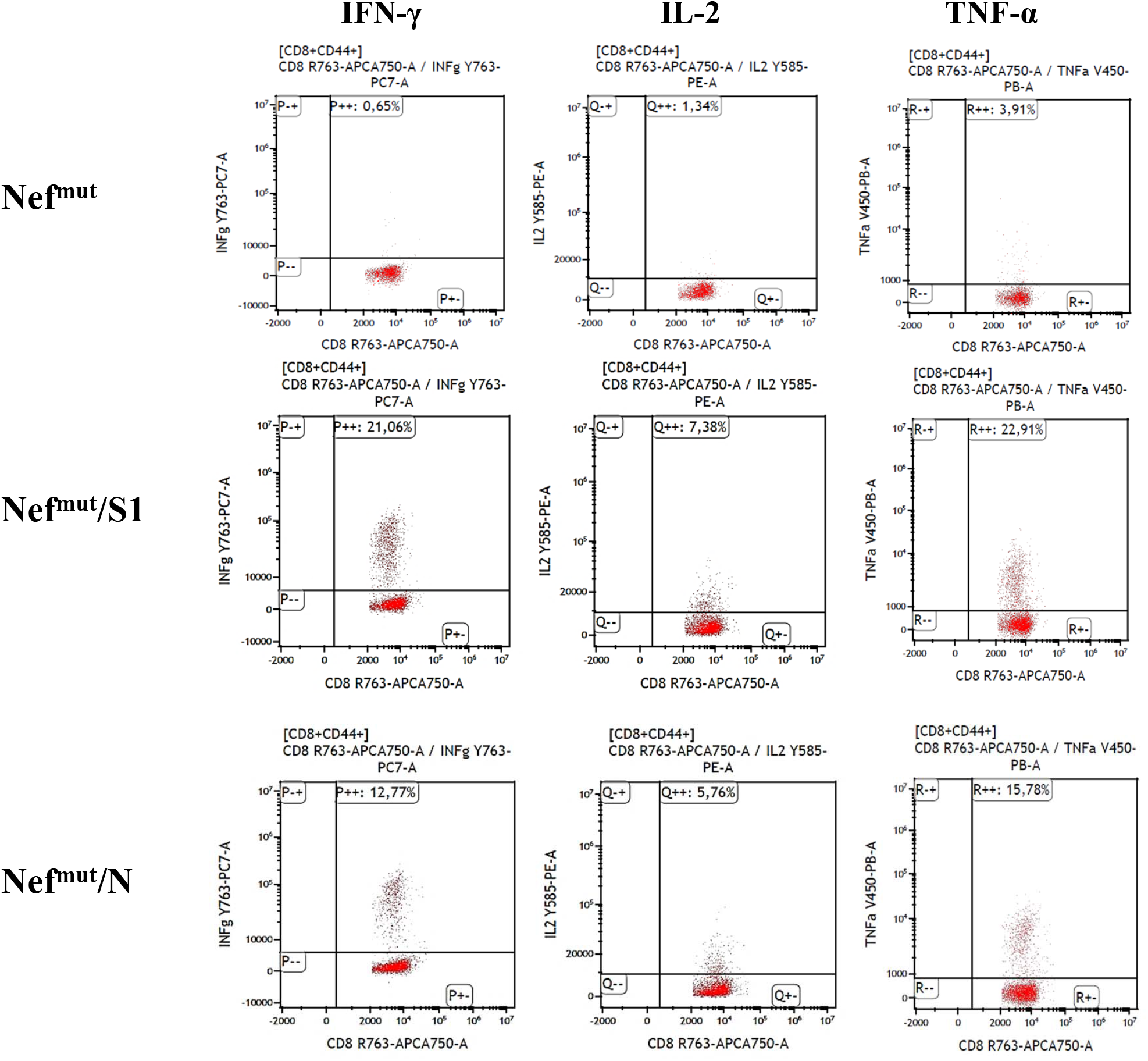
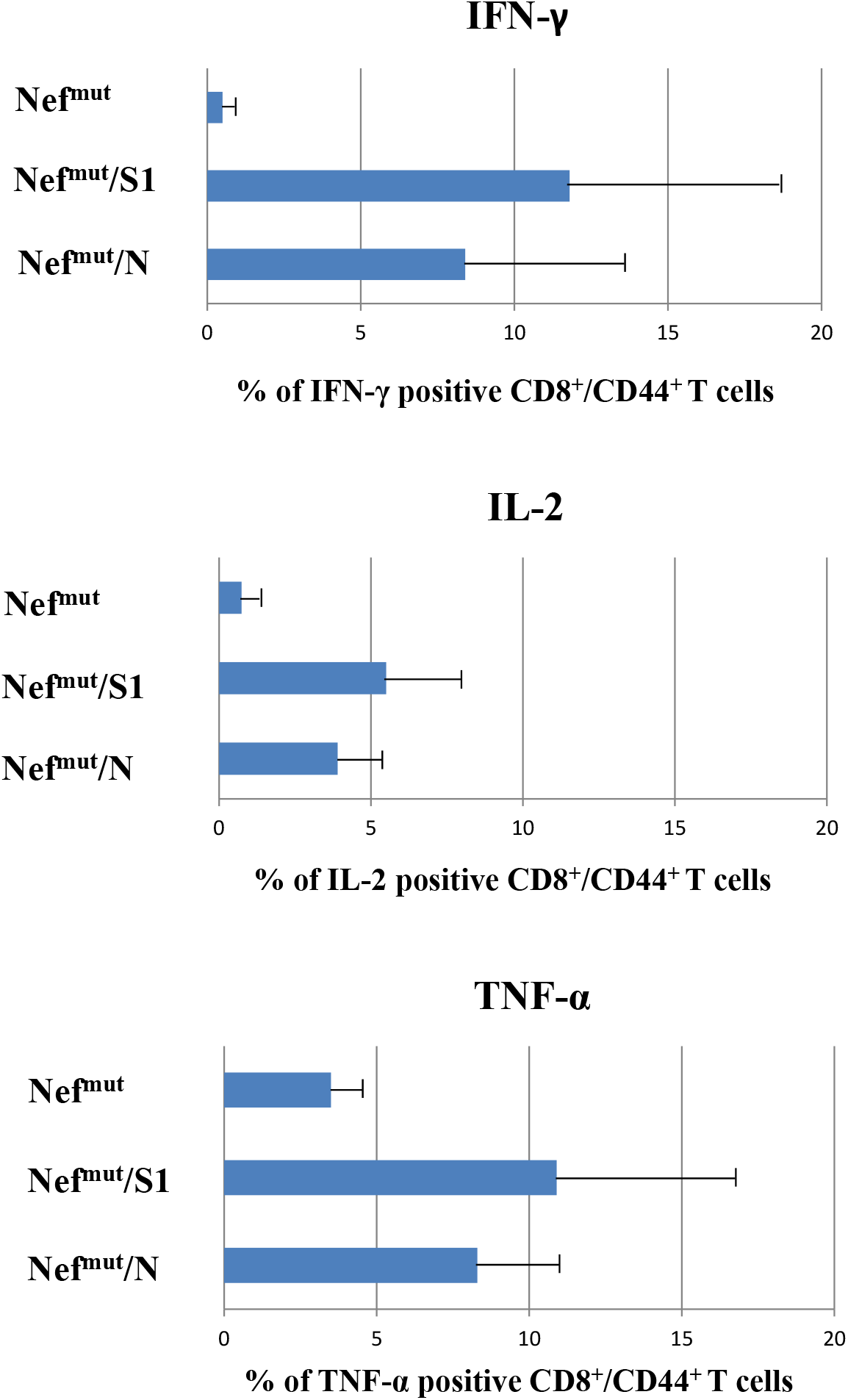
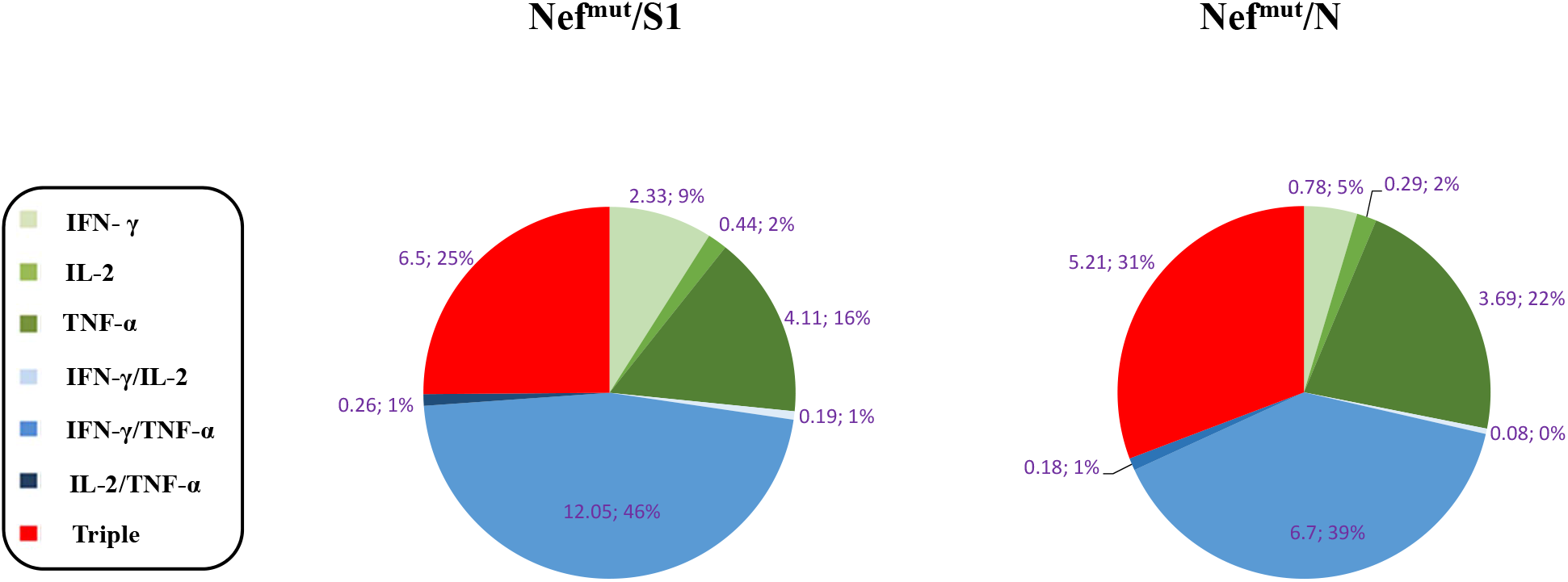
ICS/flow cytometry analysis of splenocytes from mice injected with vectors expressing either Nef^mut^/S1, Nef^mut^/N or, as control, Nef^mut^ alone. (**A**) Rough data from the analysis of the expression of IFN-γ, IL-2, and TNF-α over CD8^+^/CD44^+^ cells in splenocyte cultures from a representative mouse per group. (**B**) Percentages of cells expressing IFN-γ, IL-2, and TNF-α over the total of CD8^+^/CD44^+^ T cells within splenocytes isolated from each mouse injected with the indicated DNA vectors. Shown are mean values +SD of the absolute percentages of cytokine expressing cells from cultures treated with specific peptides after subtraction of values measured in cells treated with an unrelated peptide. The results are representative of seven independent experiments. (**C**) Pie charts reporting both absolute (i.e., over the total of analyzed CD8^+^/CD44^+^ T cells) and relative percentages of cells expressing each cytokine combination in splenocyte cultures from representative mice injected with the indicated vectors. Percentages were calculated after subtraction of values measured in homologous cultures treated with unrelated peptides.

We concluded that the injection of vectors expressing SARS-CoV-2 S1- and N-based fusion products elicited high levels of antigen-specific polyfunctional CD8^+^ T lymphocytes.

### Detection of polyfunctional antigen-specific CD8^+^ T lymphocytes in BALFs from immunized mice

Resident CD8^+^ T cells in lungs are essentially generated in an independent way respect to the pool of circulating CD8^+^ T cells, and are maintained by homeostatic proliferation to replenish the continuous loss of cells through intraepithelial migration towards lung airways [24]. Hence, a vaccine injected distally (e.g., into quadriceps), and conceived to elicit cell immunity against infectious respiratory diseases should be specifically tested for its ability to generate effective virus-specific CD8^+^ T cells in lungs. To assess the actual SARS-CoV-2-specific CD8^+^ T cell immune response in lungs after immunization with Nef^mut^-based products, cells from BALFs of at least three injected mice per group were pooled, cultivated in the presence of either specific or unrelated peptides, and analyzed for the expression of IFN-γ, IL-2, and TNF-α. Data from ICS/flow cytometry analysis showed the presence of substantial percentages of cells expressing each cytokine within CD8^+^/CD44^+^ cells isolated from BALFs (Fig. 3A), including a remarkable fraction of polyfunctional CD8^+^ T lymphocytes (Fig. 3B).

**Figure 3.**
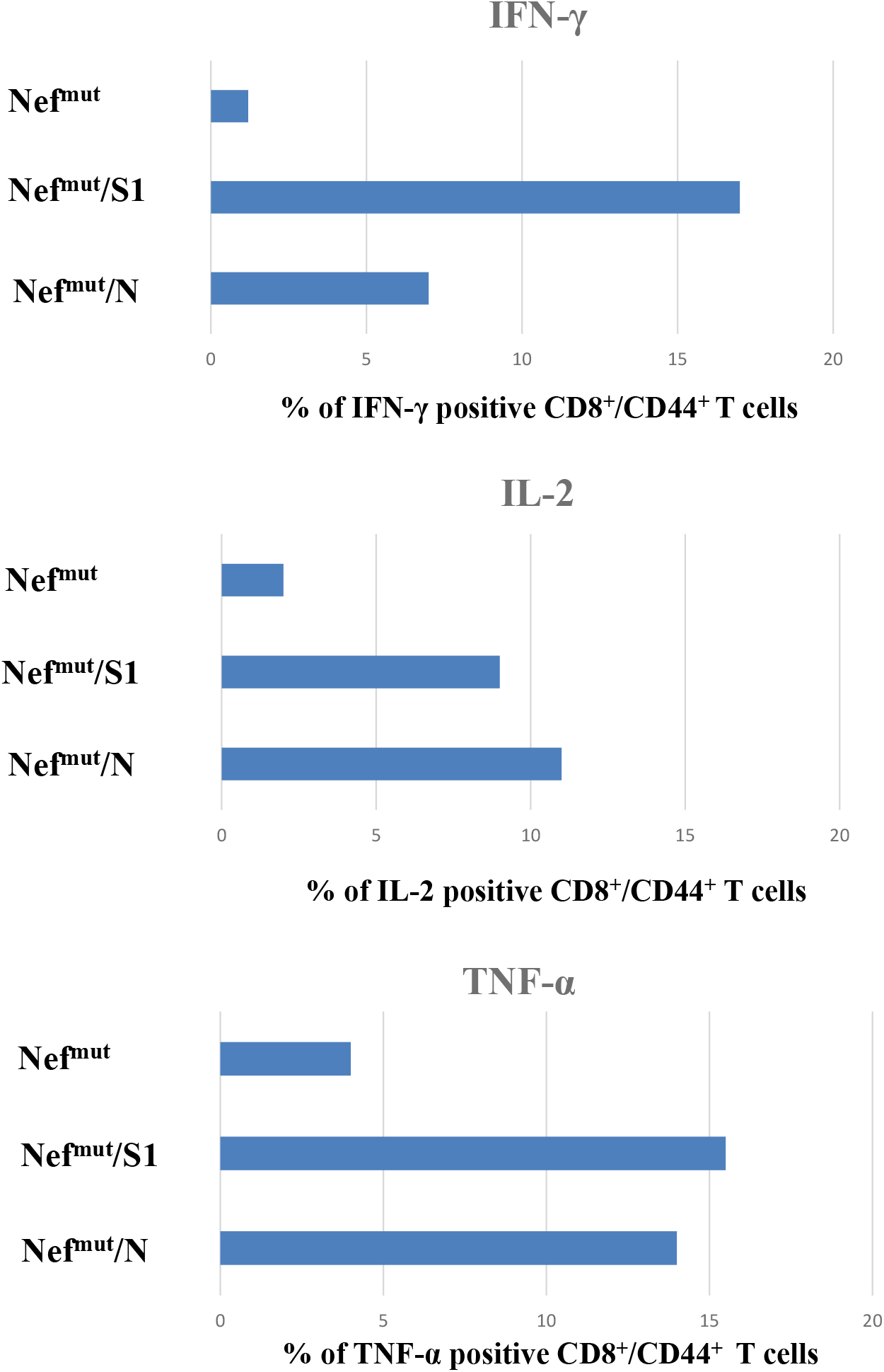
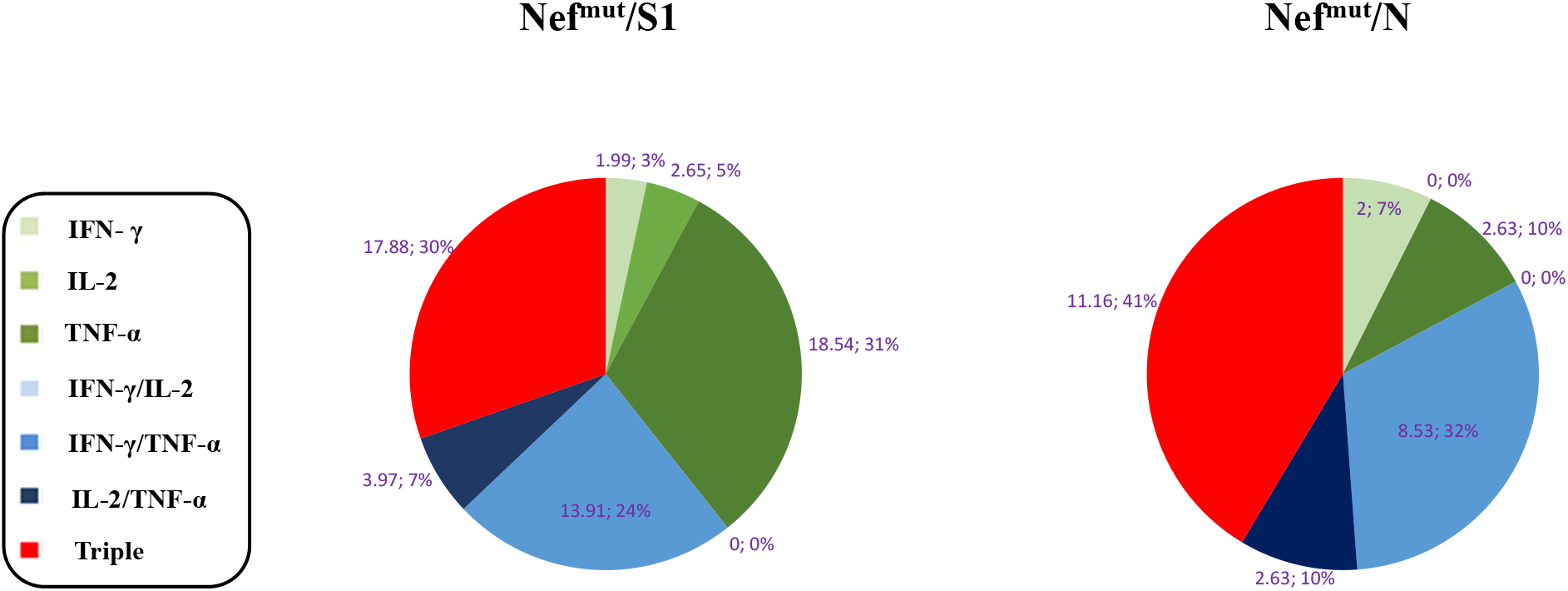
ICS/flow cytometry analysis of cells isolated from BALFs of mice injected with vectors expressing either Nef^mut^/S1, Nef^mut^/N or, as a control, Nef^mut^ alone. (**A**) Percentages of cells expressing IFN-γ, IL-2, and TNF-α over the total of CD8^+^/CD44^+^ T cells within cells pooled from at least three mice injected with the indicated DNA vectors. Shown are mean values of the absolute percentages of cytokine expressing cells from cultures treated with specific peptides after subtraction of values detected in cells treated with an unrelated peptide. The results are from two independent experiments. (**B**) Pie charts indicating both absolute (i.e., over the total of CD8^+^/CD44^+^ T cells) and relative percentages of cells expressing each cytokine combination in cells from BALFs of mice injected with the indicated vectors. Percentages were calculated after subtraction of values detected in homologous cultures treated with unrelated peptides.

This finding supports the idea that Nef^mut^-based immunization generates effective antigen-specific CD8^+^ T cells in lungs.

### Association of high levels of circulating N-specific CD8^+^ T cells with resistance to lethal SARS-CoV-2 infection

The antiviral efficacy of CD8^+^ T cell immune responses elicited against S1 and N was tested in C57 Bl/6 K18-hACE-2 transgenic mice. In these animals, the human receptor of SARS-CoV-2 virus, i.e., human angiotensin-converting enzyme (hACE)-2, is expressed under the control of the cytokeratin-18 promoter [25]. First, we proved that K18-hACE-2 transgenic mice responded similarly to the parental C57 Bl/6 strain in terms of CD8^+^ T cell immune response after injection of either Nef^mut^/S1 or Nef^mut^/N expressing vectors (Supplementary fig. 2). Next, before challenging the immunized animals, the SARS-CoV-2 preparation was titrated in terms of LD_50_. To this end, 5-fold dilutions of supernatants from infected cells, i.e., from 2.2×10^5^ to 1.8×10^3^ TCID_50_ (as measured by *in vitro* titration), were used to challenge groups of four animals (Fig. 4). We calculated that the LD_50_ corresponded to 1.0 × 10^4^ TCID_50_.

**Figure 4.**
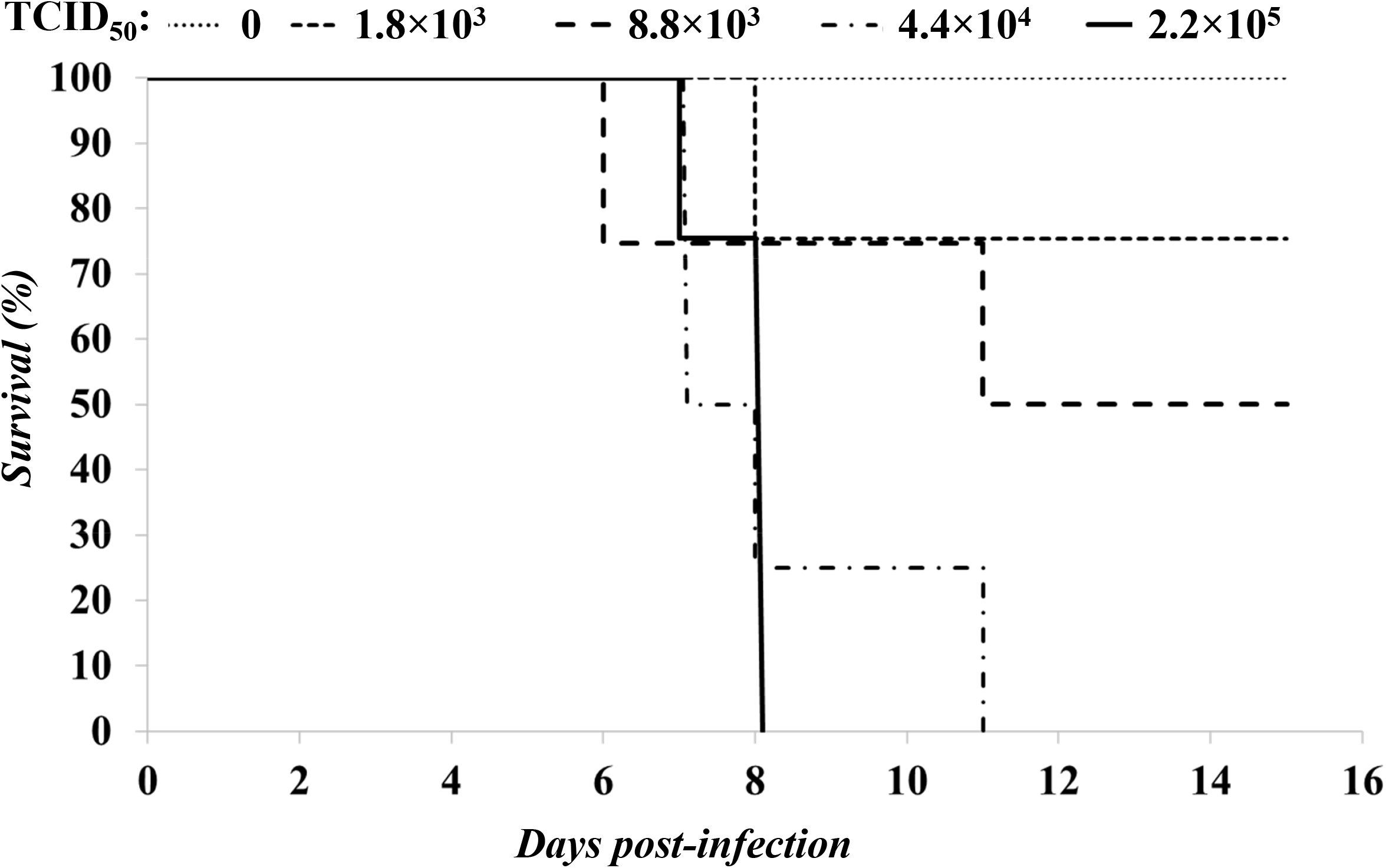
Kaplan–Meier survival curve calculated for groups (n=4) of C57 Bl/6 K18-hACE-2 mice infected with 5-fold dilutions of a SARS-CoV-2 preparation.

Three days before the infection of vaccinated mice, CD8^+^ T cell immune responses were evaluated by IFN-γ EliSpot assay using PBMCs isolated from each mouse (Fig. 5A). Then, mice were infected with 4.4 LD_50_ of SARS-CoV-2, and both weight/clinical signs of disease and survival were checked over time. In these experimental conditions, S1-immunized mice appeared as susceptible to the virus challenge as the controls (Fig. 5B-C). On the other hand, mice developing the highest levels of N-specific CD8^+^ T cell immunity resisted the lethal effect of the infection. In two of these cases, neither weight loss nor other signs of the disease (e.g., reduced reactivity) became apparent. Differences in terms of anti-N CD8^+^ T cell response levels between high and low responders were statistically significant (*p* = 0.0286, one-tailed Mann-Whitney U test). Consistently, the highest levels of N-specific CD8^+^ T cells associated with protection in a group of mice contemporarily injected with vectors expressing Nef^mut^/S1, Nef^mut^/S2, and Nef^mut^/N (Supplementary fig. 3). Conversely, in both instances, mice with low counts of N-specific CD8^+^ T cells succumbed.

**Figure 5.**
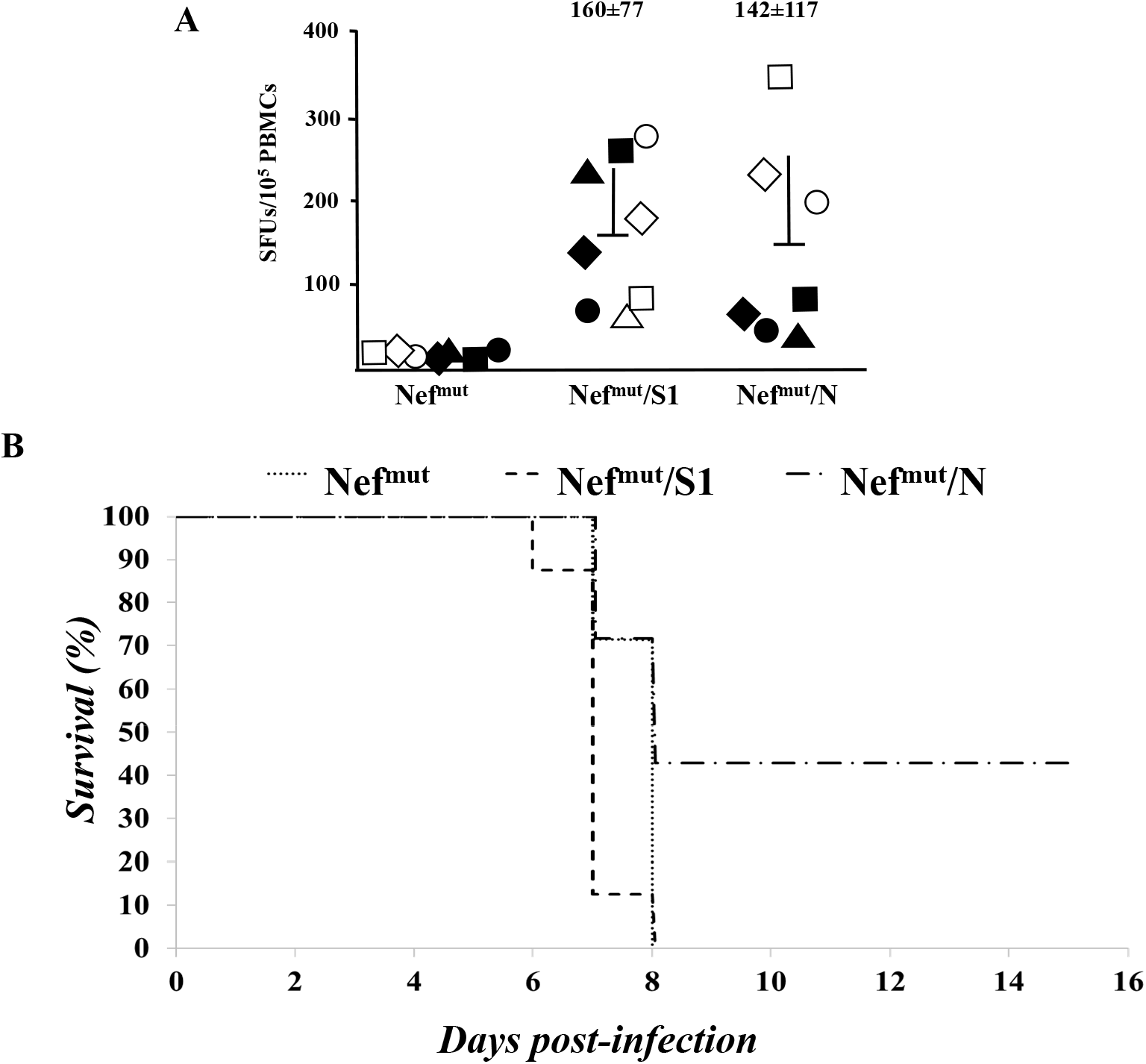
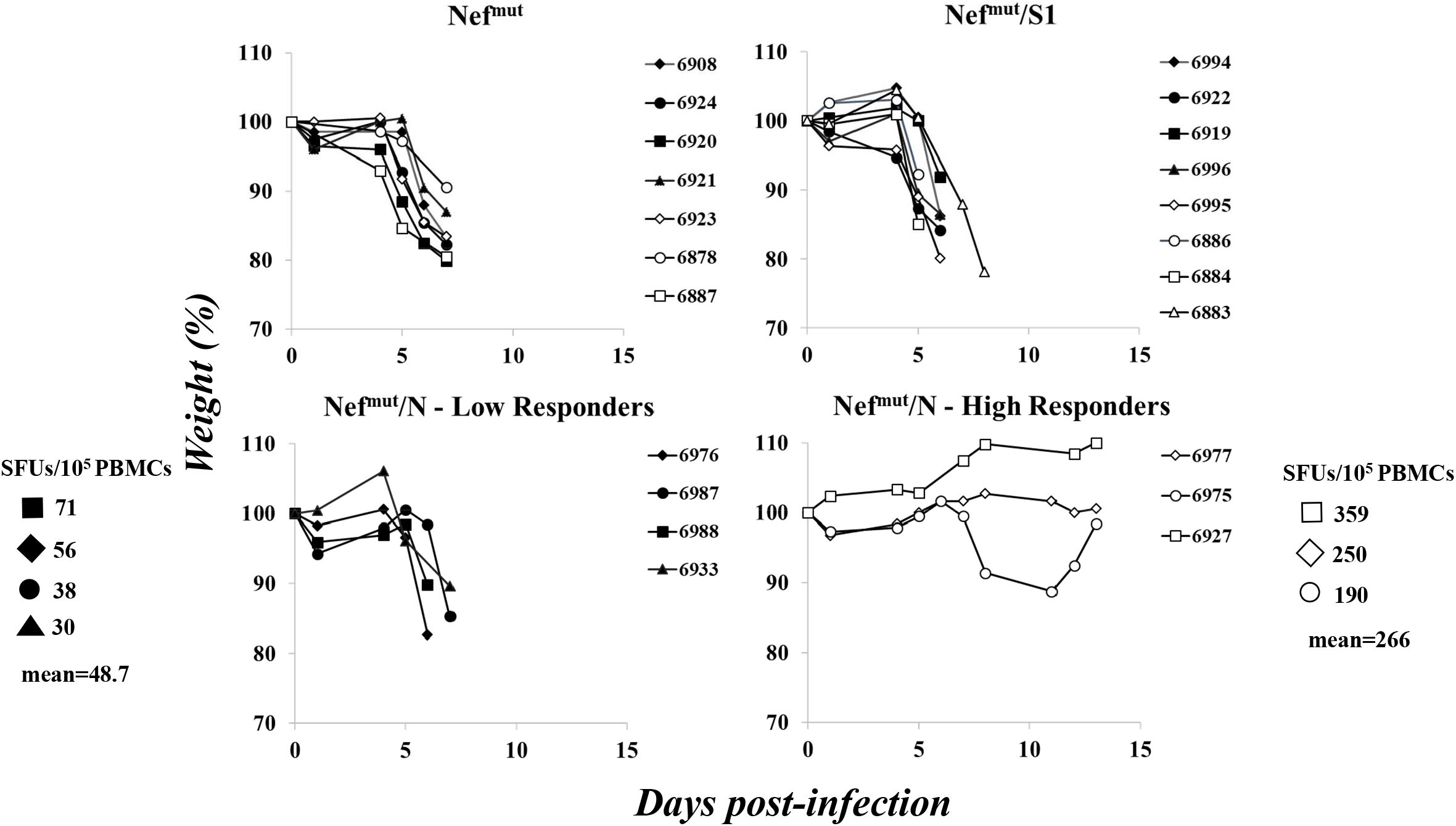
Antiviral effect induced by high levels of N-specific CD8^+^ T cell immunity. (**A**) CD8^+^ T cell immune response in C57 Bl/6 K18-hACE-2 mice injected with vectors expressing either Nef^mut^/S1, Nef^mut^/N or, as control, Nef^mut^ alone. PBMCs were isolated by retro orbital bleeding and, after erythrocyte lysis, were incubated o.n. with or without 5 μg/mL of either unrelated or SARS-CoV-2 related peptides in triplicate IFN-γ EliSpot microwells. Shown are the number of SFUs/10^5^ PBMCs as mean values of triplicates after subtraction of values from wells treated with an unrelated peptide. Intragroup mean values + SD are reported. (**B**) Kaplan–Meier survival curve calculated for groups of C57 Bl/6 K18-hACE-2 mice infected with 4.4 LD_50_ of SARS-CoV-2. Differences between Kaplan-Meier survival curves relative to S1- and N-immunized groups of mice were statistically significant (log-rank test, *p* = 0.01285). (**C**) Relative weight loss in each injected mice after SARS-CoV-2 challenge. Identification numbers of each mouse are reported on the right of each panel. SFUs/10^5^ PBMCs for each low and high responder N-immunized mouse are also indicated together with intergroup mean values. Shown are cumulative data from two experiments.

Taken together, these data indicated that adequate levels of CD8^+^ T cell immunization against SARS-CoV-2 N confer resistance against the lethal effect of SARS-CoV-2.

### Detection of N-specific CD8^+^ T-resident memory (Trm) cells in lungs of immunized mice

The overall quality of any vaccine strategy relies also on the duration of the protective effect. The kinetics of protection, in turn, strictly depends on generation of antigen-specific memory cells. In the case of CD8^+^ T cell-based immunity against respiratory viruses, the induction of virus specific CD8^+^ Trm cells in lungs is mandatory to ensure a reliable duration of the immunity. We tried to assess whether the i.m. injection of Nef^mut^/N expressing vector was sufficient to induce N-specific CD8^+^ Trm cells in lungs. To this end, immune cells were isolated from lungs of mice injected with either Nef^mut^/N or control vectors. After overnight treatment with either N-specific or unrelated peptides, the CD8^+^/CD44^+^ sub-populations were scored by ICS/flow cytometry analysis for the simultaneous expression of IFN-γ and Trm cell markers, i.e., CD49a, CD69, and CD103 (Fig. 6A). Through Boolean-gating based analysis, we reproducibly identified an N-specific, CD8^+^ Trm cell sub-population in lungs of mice injected with the Nef^mut^/N expressing vector (Fig. 6B).

**Figure 6.**
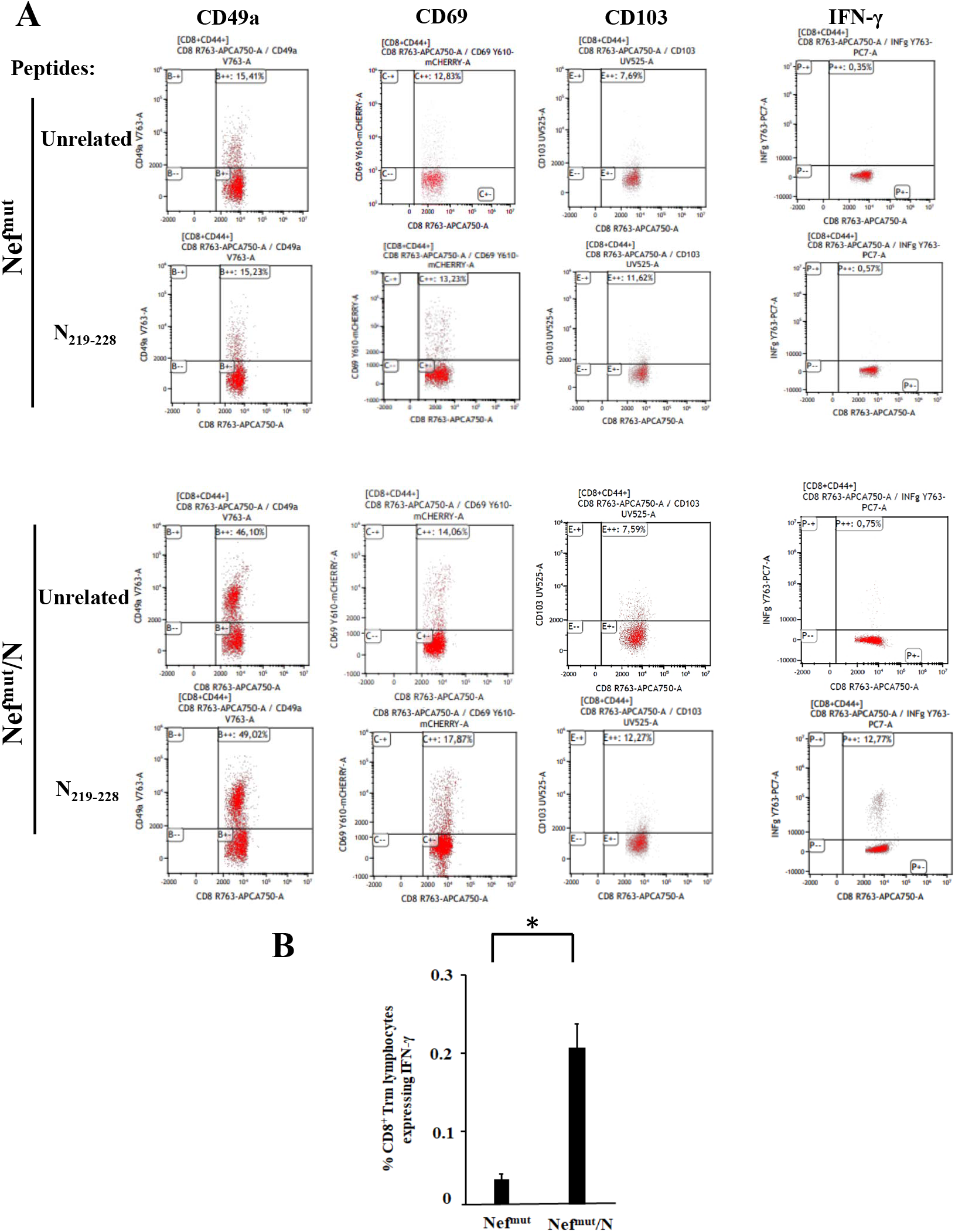
ICS/flow cytometry analysis on cells isolated from lungs of mice injected with vectors expressing either Nef^mut^/N or Nef^mut^ alone. (**A**) Rough data obtained by analyzing cells pooled from lungs of two representative mice per group. (**B**) Percentages of CD8^+^ Trm cells expressing IFN-γ over the total of CD8^+^/CD44^+^ T lymphocytes. Shown are mean values + SD of the absolute percentages of positive cells from cultures treated with specific peptides after subtraction of values detected in cells treated with an unrelated peptide. The results are representative of three independent experiments. \**p*<0.05.

We concluded that the N-specific immunity conferred by the injection of Nef^mut^/N-expressing vector can lead, besides protection against lethal virus infection, to the establishment of CD8^+^ T memory cells at lungs. This finding could be of outstanding relevance in the perspective of a translation into the clinic of the Nef^mut^-based vaccine platform.

## DISCUSSION

HIV-1 Nef^mut^ is a protein mutant lacking the pathogenic effects induced by the wild-type counterpart [26]. It incorporates into nascent EVs at quite elevated extents also when heterologous proteins are fused at its C-terminus [16]. When DNA vectors expressing Nef^mut^-derivatives are i.m. injected, EVs spontaneously released by muscle cells upload Nef^mut^-based products [22]. Engineered EVs can freely circulate into the body, and their internalization into APCs generates a CTL immunity against EV-uploaded antigens in the absence of induction of specific antibodies [17]. The CTL immune response depends on both amounts and efficiency of DNA delivery, lasts several months in peripheral circulation and spleen, and protects mice from both HPV16- and HER2-related cancers [12, 13]. The strength of CD8^+^ T immunity elicited by expression of antigens fused with Nef^mut^ largely exceeds that induced by the expression of foreign antigens alone [12, 23].

The Nef^mut^-based method of CD8^+^ T cell immunization was found effective also when applied on SARS-CoV-2 antigens [18]. However, the actual efficacy of this strategy strictly depends on the generation of effective cell immunity at lungs. In fact, in view of the documented compartmentalization of both B- and T-cell immunity in lungs [24, 27], the appearance of antigen-specific CD8^+^ T cells in peripheral circulation/secondary lymphoid organs does not necessarily imply a contemporary induction of cell immunity in airway tissues. On this subject, the here described detection of SARS-CoV-2 specific polyfunctional CD8^+^ T cells in BALFs from injected mice demonstrated the effectiveness of the Nef^mut^- based method in establishing a CD8^+^ T cell immunity in lungs.

In view of the compartmentalization of lung cell immunity, how can be it reconciled the prompt induction of lung cell immunity with the administration of the immunogen into a distal district, i.e. quadriceps? We hypothesize that the immunity we found in airways was consequence of intra-tissue diffusion of engineered EVs. In support of this idea, data from several reports demonstrated that lungs, together with spleen, liver and bone marrow, can be populated by fluorescently labelled EVs shortly after intravenous injection [28-31]. Similarly, Nef^mut^-engineered EVs emerging from muscle cells can access airway districts, where they can be internalized by resident APCs. Thereby, EV-uploaded products can be cross-presented, and CD8^+^ T lymphocyte selection and activation can be initiated in the context of local germinal centers and/or mediastinal lymph nodes. We assumed that the levels of antigen-specific CD8^+^ T lymphocytes in both peripheral circulation and lungs were directly proportional to the amounts of produced engineered EVs. In such a scenario, predicting the SARS-CoV-2-specific immunologic status in lungs by measuring the related CD8^+^ T cell immunity in spleen and peripheral circulation seems appropriate.

In our experiments, S1- and N-specific CD8^+^ T cell immunity seemed both quantitatively and qualitatively comparable. However, protection from lethal SARS-CoV-2 challenge was observed only in mice developing the highest N-specific immune responses. The lack of protection in mice less efficiently immunized against N can be readily interpreted in terms of a quantitatively inadequate immune response. Conversely, the interpretation regarding the lack of virus containment by S1-specific CD8^+^ T cell immune response is not obvious. The apparently lower levels of S1 uploading in EVs compared to N [18] should not be a critical issue, since the S1-specific immune response appeared similar or even slightly stronger than that induced by N in nearly all tests we carried out. Conversely, the inefficacy of S1-specific immunity might be consequence of low accumulation of SARS-CoV-2 S in live infected cells, due to its spontaneous association with cell plasma membrane and shedding. On the other hand, higher intracellular accumulation of N can result in more molecules available for degradative pathways and MHC Class I peptide presentation. A more efficient exposition of N-derived peptides would in turn result in a quicker recruiting of specific CD8^+^ T lymphocytes at the site of virus-infected cells, ultimately leading to a more efficient containment of viral spread.

Using a viral dose as high as 4.4 LD_50_, we observed protection in mice that had developed more robust N-specific CD8^+^ T cell immunity. The results in terms of protection from virus challenge seemed to be influenced by some variability in the levels of immunity reached after two injections, similarly to what previously observed in mice immunized by EVs engineered by Nef^mut^ fused with HPV16-E6 and -E7 [12]. Also in this case, mice resisted the tumor implantation carried out before vaccination only in the presence of adequate E6- and E7-specific CD8^+^ T cell immune responses.

The induction of N-specific CD8^+^ Trm cells in lungs represents a relevant value added for the Nef^mut^- based vaccine strategy against SARS-CoV-2. Considering that the expression of both CD49a and CD103 markers on CD8^+^ T lymphocytes has been associated with cytotoxic functions [32-34], our findings support the idea that the Nef^mut^-based immunization has the potential to generate a long lasting, effective antiviral immunity in lungs.

Our study presents limitations. The use of fluorescent tetramers in ICS/flow cytometry analyses would have identified SARS-CoV-2-specific CD8^+^ T cell sub-populations more accurately. In addition, challenging immunized mice with sub-lethal viral doses may have revealed potentially protective effects induced by S1-specific CD8^+^ T cells also. Furthermore, the lack of quantification of viral loads in lungs of protected mice precluded the evaluation of the effects of N-specific immunity on viral spread within airway tissues.

Emergence of variants against which anti-S neutralizing antibodies lose potency represents one of the most relevant shortcoming for current anti-SARS-CoV-2 vaccines. Notably, N-specific antiviral CD8^+^ T cell immunity is not expected to suffer from such a limitation. In fact, as reported in fig. 7, amino acid sequences of N protein from current variants of concern (VOCs) are well conserved, at least in part because mutations in this key viral component could have detrimental effects on optimal viral fitness. By consequence, the N-specific CD8^+^ T cell immune response against a single viral strain is anticipated to be effective also against other VOCs. In addition, based on the observations on patients recovered from SARS-CoV infection [11], SARS-CoV-2 N-specific CD8^+^ T cell immunity would wane with a kinetics much slower than that of anti-S neutralizing antibodies.

**Figure 7.**
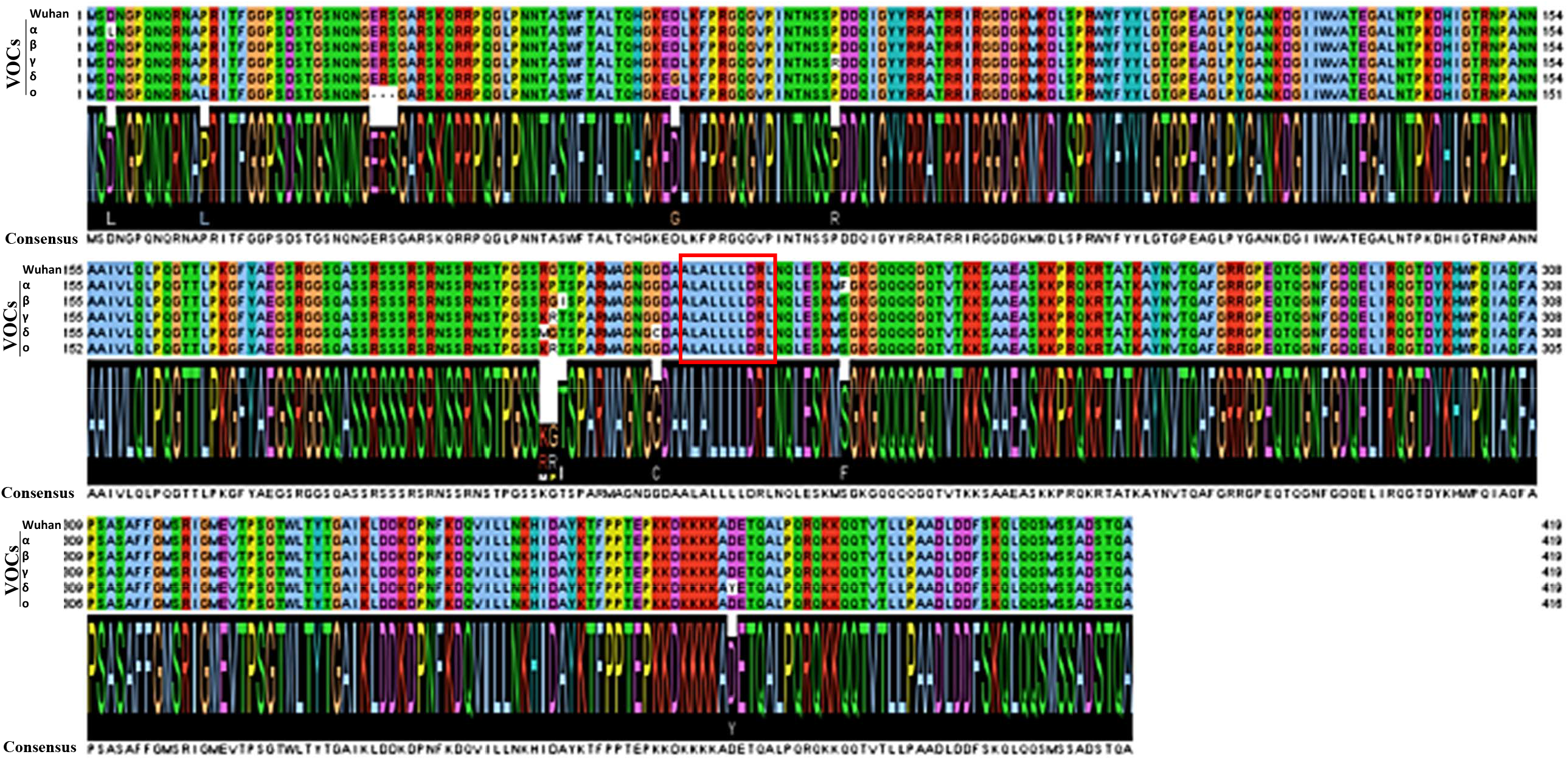
Line up of amino acid sequences of SARS-CoV-2 N protein from ancestral (Wuhan) and a number of variants of concern (VOCs). Sequences of the highly conserved H2-b immunodominant N_219-228_ epitope are highlighted.

We here demonstrate that adequate levels of N-specific CD8^+^ T cell immunity correlate with protection against lethal SARS-CoV-2 infection. As recently outlined in a seminal paper, SARS-CoV-2-specific CD8^+^ T cell immunity can be fully protective in the context of not adequate levels of neutralizing antibodies [2]. On this basis, a combination mRNA-based vaccine strategy designed to contemporarily induce N-specific CD8^+^ T immunity through engineered EVs and anti-S neutralizing antibodies is expected to overcome the limitations of current vaccines in terms of diminished efficiency against VOCs [35] and immunity waning [36]. In addition, here presented data open the way towards the exploitation of the Nef^mut^-based vaccine platform in the fight against additional respiratory viruses, like Influenza and Respiratory Syncytial viruses.

## Acknowledgments

Both SARS-Related Coronavirus 2, Isolate Italy-INMI1, NR-52284, and the peptide array of SARS-Related Coronavirus 2 Spike Glycoprotein, NR-52402, were obtained through BEI Resources, NIAID, NIH. We thank Michele Equestre, Istituto Superiore di Sanità, for deep sequencing the SARS-CoV-2 isolate. We also thank Pietro Arciero Istituto Superiore di Sanità, for technical support, and Federica Magnani and Rosangela Duranti Istituto Superiore di Sanità for secretarial assistance.

## LEGEND TO FIGURES

**Supplementary fig. 1.**
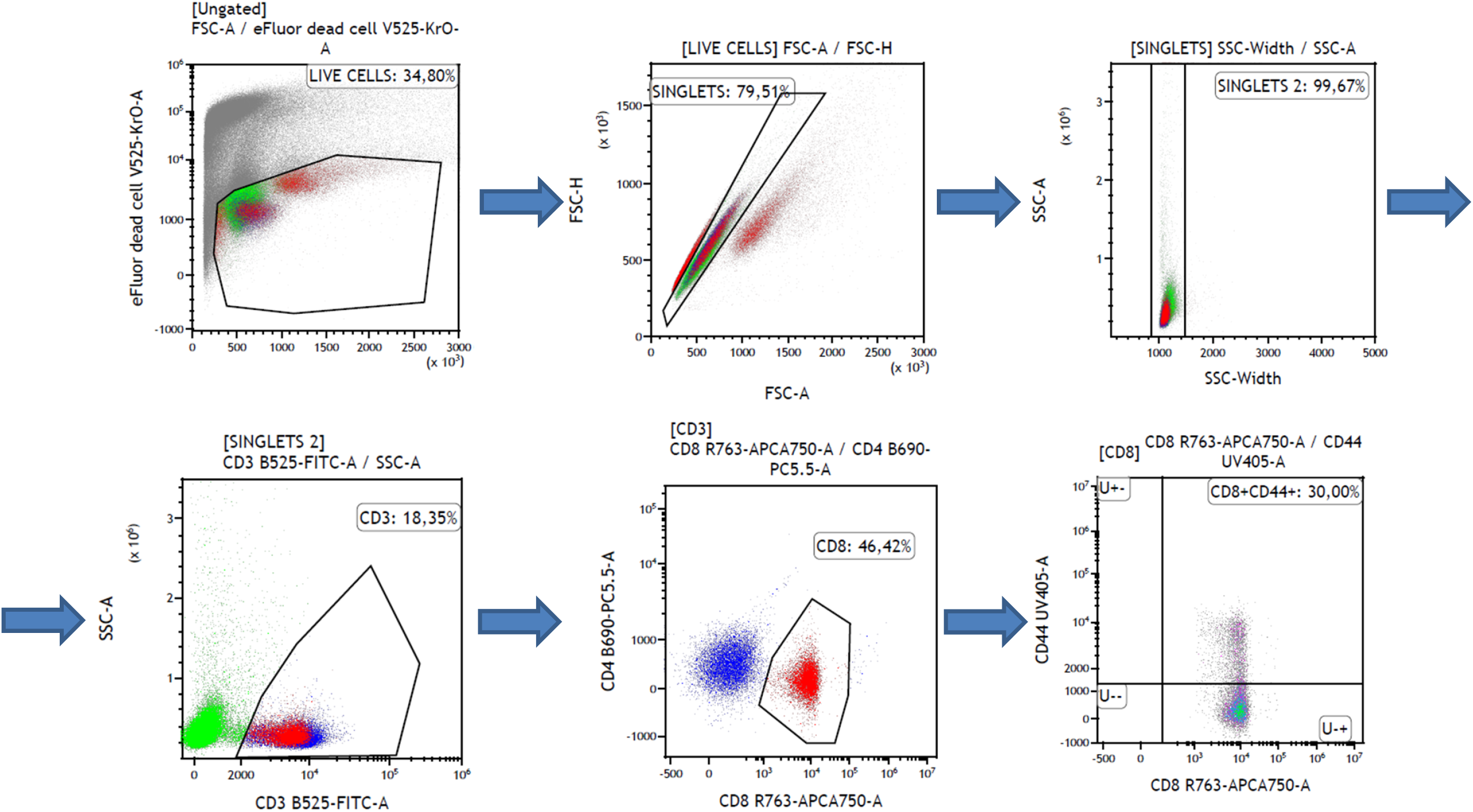
Gating strategy carried out in ICS/flow cytometry analysis of splenocytes from injected mice. Shown is the analysis on PMA-treated cells.

**Supplementary fig. 2.**
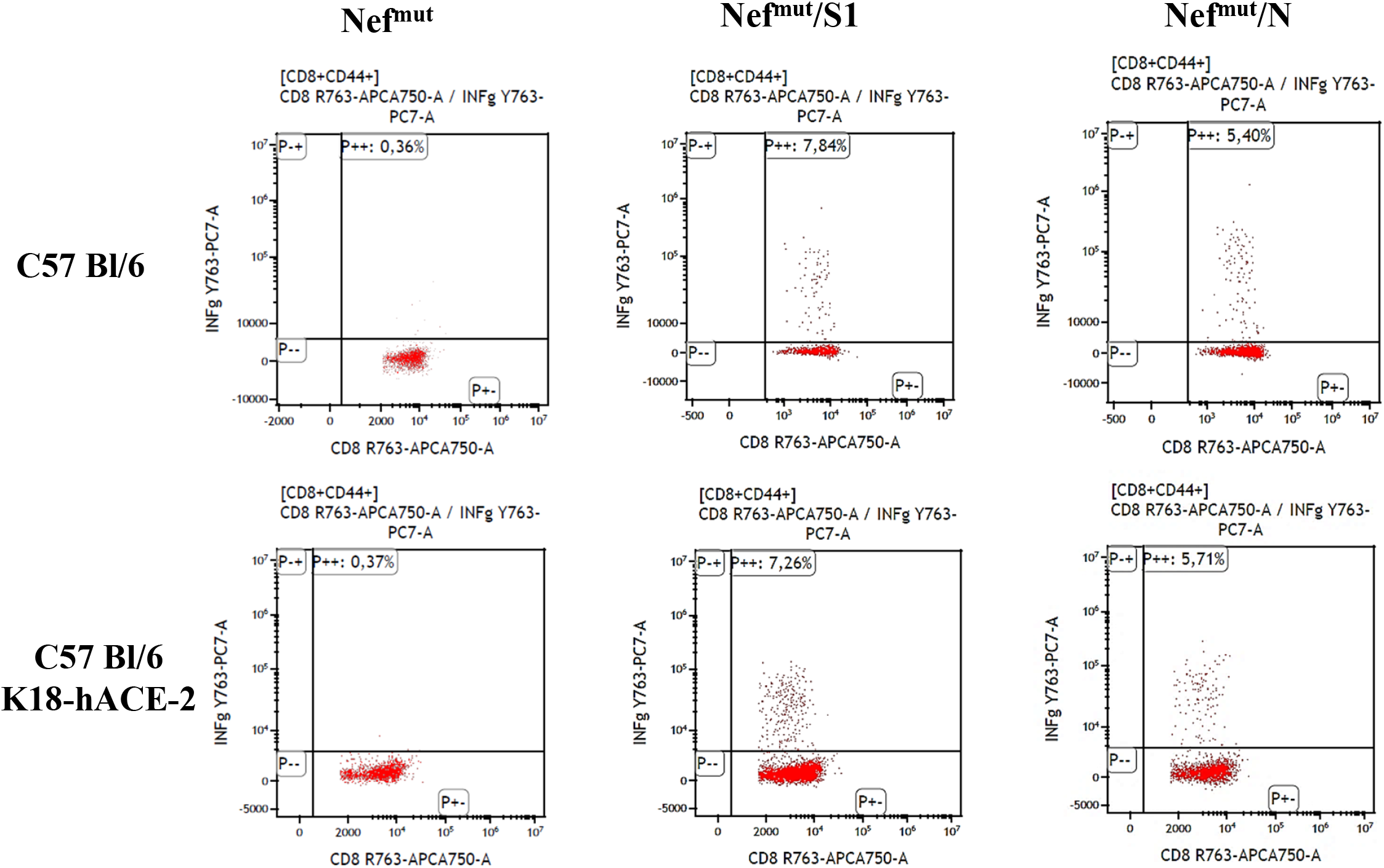
ICS/flow cytometry analysis for the expression of IFN-γ in splenocytes from either C57 Bl/ or K18-hACE-2 mice injected with vectors expressing either Nef^mut^/S1, Nef^mut^/N or, as control, Nef^mut^ alone, and cultivated overnight with SARS-CoV-2 specific peptides. Shown are rough data representative of the results obtained with splenocytes pooled from two mice per condition. Quadrants were set on the basis of cell fluorescence of samples treated with an unrelated peptide.

**Supplementary fig. 3.**
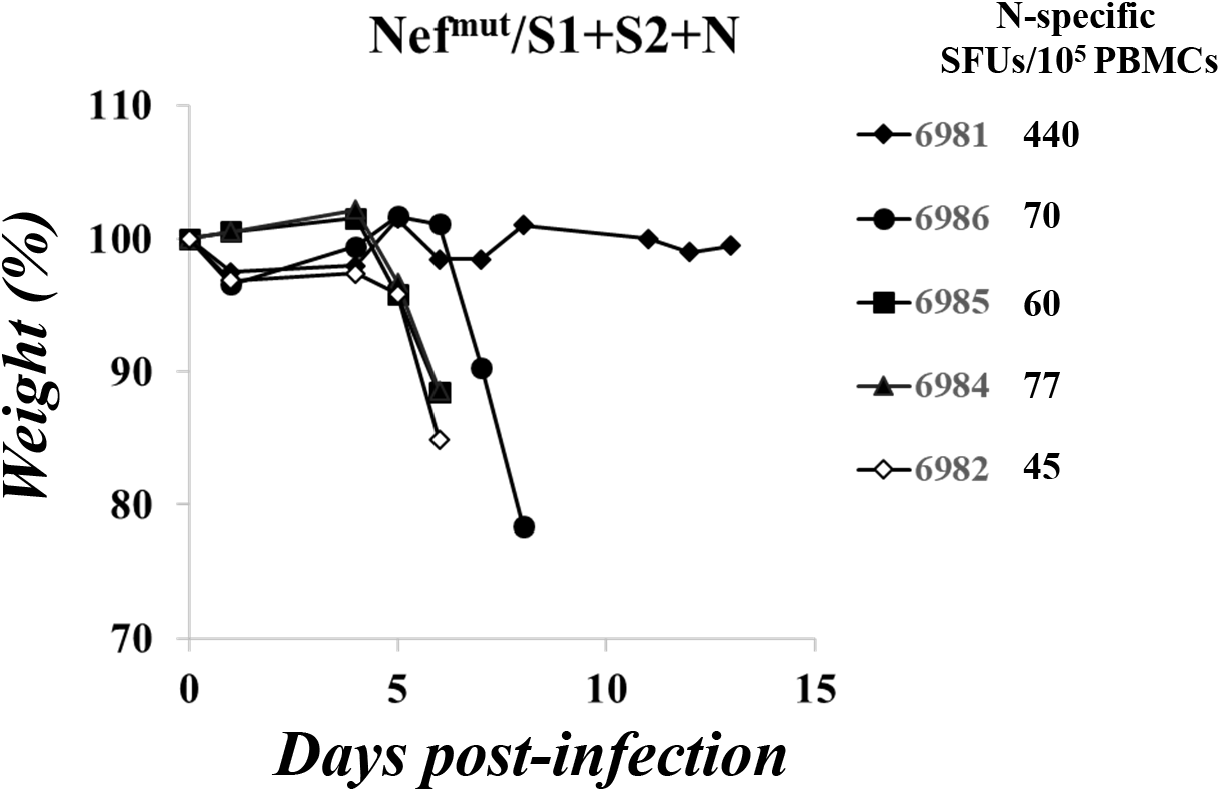
Relative weight loss in mice injected with 10 μg each of Nef^mut^/S1, Nef^mut^/S2 and Nef^mut^/N expressing vectors, and then infected with 5 LD_50_ of SARS-CoV-2. Identification numbers for each mouse are reported on the right together with the counts of SFUs/10^5^ PBMCs after incubation with the N_219-228_ peptide. S2-specific SFU counts as measured using a pool of S2 peptides were similar in each mouse.

